# *Cis*-regulation of gene expression between sexes and life stages in *Rumex hastatulus*

**DOI:** 10.1101/2025.06.16.659834

**Authors:** Meng Yuan, Bianca M. Sacchi, Baharul I. Choudhury, Spencer C. H. Barrett, John R. Stinchcombe, Stephen I. Wright

## Abstract

The potential for conflict between sexes and life stages while sharing predominantly the same genome has important evolutionary consequences. In dioecious angiosperms, genes beneficial for the haploid pollen stage may reduce the fitness of diploid offspring of both males and females. However, we still lack an understanding of the extent of shared genetic architecture for gene expression between the sexes or life stages in plants, a key component for predicting the potential for conflict. We performed expression quantitative trait loci (eQTL) mapping to test if standing variation affects sexes and life stages differently using a population sample of the dioecious outcrossing plant *Rumex hastatulus*. We compared effect sizes and allele frequencies of *cis*-eQTLs in male and female leaf tissues and pollen and tested for genotype-by-sex interactions for gene expression. We found stronger shared genetic architecture between sexes than between life stages, suggesting greater potential for ongoing sexual conflict in leaves, which have been shown to be sexually dimorphic in earlier studies. In contrast, conflict over optimal gene expression between pollen and leaves may be easily resolved due to their distinct genetic architectures. Additionally, our burden of rare allele test suggested a signature of stabilizing selection against extreme gene expression in leaves. Our study highlights the use of eQTLs to investigate selection on gene expression and the evolution of conflict between sexes and life stages in dioecious species.

## Introduction

Evolutionary conflict may arise when different sexes or life-cycle phases of an organism have different fitness optima while sharing a common genome (Ewing 1977; Arnqvist and Rowe 2005; Haig and Wilczek 2006; Schärer et al. 2015). For example, sexual conflict occurs when males and females in dioecious species have different fitness optima involving reproduction, and this can drive sexual dimorphism in phenotypic traits (Lande 1980; Bonduriansky and Chenoweth 2009). In plants, the diploid and haploid phases of the life cycle (i.e., sporophytes and gametophytes respectively, we use “life stages” to refer to these phases) are distinct in their morphology, physiology, and development despite sharing extensive overlap in gene expression, creating potential for genetic trade-offs (Haig and Wilczek 2006; Beaudry et al. 2020). Both sexual and life-stage conflict have been suggested to generate balancing selection and stable genetic variation under some conditions (Kidwell et al. 1977; Immler et al. 2012). However, we lack comprehensive understanding of the traits and genetic architecture underlying the conflict between sexes or life stages. Here, we use expression quantitative trait locus (eQTL) mapping to identify genome-wide *cis*-regulatory variation and investigate the extent of shared genetic architecture between sexes and life stages and the potential for sexual and life-stage conflict in a dioecious plant.

Current empirical examples of life-stage conflict often involve both sexes (i.e., a conflict both between the sexes and life stages). For example, alleles beneficial for haploid gametophytic competition were shown to reduce the fitness of female diploid offspring in the hermaphroditic annuals *Clarkia* (Travers and Mazer 2001) and *Collinsia* (Lankinen and Kiboi 2007; Lankinen et al. 2017). Sexual conflict independent of life stage differences has been observed in several plant species (e.g., Delph et al. 2011; Duffy et al. 2021; Wang et al. 2021), but genomic tests of sexual conflict in plants remain limited compared to animals (but see Delph et al. 2022). Gene expression is an important molecular trait underlying phenotypic variation. While direct tests of conflict require measurements of fitness which is often laborious and challenging, sexual or life-stage dimorphism in gene expression has been widely used to indirectly study conflict (Mank 2017; Yuan et al. 2025). With the considerable potential for both sexual and life-stage conflict in plants, whether there is a greater likelihood for one type of conflict than the other, or whether one type of conflict is more resolved than the other, remains unanswered. Thus, plants offer excellent systems to study the evolution of conflict between sexes and life-cycle phases in a single species.

The potential for future conflict, i.e., whether new mutations and standing variation are more likely to be pleiotropic, depends on the degree of shared genetic architecture underlying phenotypic traits between sexes or life stages. Genetic architecture describes how genetic variation shapes phenotypic differences, including the number and effect sizes of contributing genes. When both sexes share most of the genetic architecture underlying a trait (i.e., a positive and high intersexual genetic correlation *r*_mf_), there is strong potential for ongoing conflict in the presence of antagonistic selection, and the evolution of sexual dimorphism is constrained (Lande 1980; Poissant et al. 2010; Wyman et al. 2013). Genetic covariances between sexes inferred through gene expression studies in *Drosophila* suggest considerable scope for ongoing conflict (Allen et al. 20182018; Houle and Cheng 2021; Singh and Agrawal 2023; Grieshop et al. 2025). In contrast, when there is distinct genetic architecture between sexes or life stages, traits can evolve towards the same optima under concordant selection, while dimorphisms can evolve under antagonistic selection. Testing the extent of shared genetic architecture genome-wide is an important first step to understand the potential for ongoing of conflict between sexes and life stages.

Expression quantitative trait loci (eQTL) mapping is useful for identifying loci underlying phenotypic variation or those subject to selection (Kudaravalli et al. 2009; Lee et al. 2014; Josephs et al. 2015). Local eQTLs, i.e., genetic variants that are significantly associated with the expression level of a closely linked focal gene, are differentiated from distant eQTLs by a distance threshold in eQTL mapping studies. Most local regulatory variation acts in *cis* and has allele-specific effects; we refer to local eQTLs as *cis*-eQTLs or *cis*-regulatory variation. *Cis*-regulatory variation has a relatively large effect on gene expression, making it easier to map *cis*-eQTLs in association studies (Josephs et al. 2017; Signor and Nuzhdin 2018). Whether *cis*-eQTLs affect different sexes or life stages concordantly can be used to infer the extent of shared genetic architecture between sexes and life stages. For example, a study on eQTLs across human tissues has shown limited standing variation with sex-specific effects (Genotype × Sex interaction) in expression, suggesting high potential for sexual conflict under sexually antagonistic selection (Oliva et al. 2020). Furthermore, selective pressure on eQTLs can be inferred from their distribution of allele frequencies when compared to the neutral expectation. In the leaf tissues of *Capsella grandiflora, cis*-eQTLs showed excessive rare alleles consistent with purifying selection (Josephs et al. 2015); whereas in floral tissues of *Mimulus guttatus, cis-* eQTLs exhibited more common than rare alleles (Brown and Kelly 2021). These studies of two plant species suggest that eQTLs can experience different selective pressures, and that potentially differential selection can act on genes expressed in different sexes and/or life stages.

We investigated *cis*-regulation of gene expression through *cis*-eQTL mapping in the dioecious, wind-pollinated and obligately outcrossing annual *Rumex hastatulus* (Polygonaceae). Several previous findings in *R. hastatulus* indicated the potential for antagonism between sex and life stages, especially during pollen competition. Sexual dimorphism in phenotypic and life history traits of this species is likely associated with differential reproductive roles between sexes during wind pollination, e.g., males are taller and produce more inflorescences during peak flowering to maximize pollen dispersal while females are taller and have more inflorescences at reproductive maturity to maximize seed dispersal (Puixeu et al. 2019). Vegetative traits such as leaf morphology are tightly linked to survival and reproductive success (e.g., Delph et al. 2011; Scharmann et al. 2021). In *R. hastatulus*, leaf size is positively correlated with plant height in both sexes, and both leaf number and size are sexually dimorphic (Puixeu et al. 2019). The high intensity of pollen competition in wind pollinated plants provides an opportunity to test gametophytic selection during pollen competition (Friedman and Barrett 2009; Field et al. 2012) and its pleiotropic effects on diploid fitness, e.g., a potential trade-off between pollen success and diploid offspring fitness. Selection during pollen competition has been shown to affect sex-ratio bias (Field et al. 2012) and sex chromosome evolution (Sandler et al. 2018) in *R. hastatulus*, e.g., female biased sex ratios are likely caused by greater competitive ability of X-bearing pollen following Y chromosome degeneration (Hough et al. 2014; Crowson et al. 2017). The potential conflict between sexes and life stages in *R. hastatulus* makes it a promising experimental system to test the direction of selection on gene expression between sexes and life stages and also to compare the potential for sexual and life-stage conflict.

We tested differential gene expression between sexes and life stages and mapped *cis*-eQTLs separately in leaf and pollen tissues to compare their effect sizes and allele frequencies. We found significantly fewer differentially expressed genes between sexes than between life stages, and most of the highly sex-biased genes were on the sex chromosomes. There was a greater overlap in eGenes (i.e., genes with their expression level affected by eQTLs) and eQTLs between sexes than life stages. Effect sizes of eQTLs were positively correlated between sexes and life stages, with a much stronger correlation between sexes, indicating more shared genetic architecture between sexes. Consistent with this result, we found limited Genotype × Sex interaction in gene expression. Lastly, we found similar selective pressures in eQTLs between sexes and life stages based on their allele frequency distributions but with no excess of rare alleles among eQTLs in any tissues when compared to a null distribution. Rare genetic variation upstream of each gene exhibited both up- and down-regulated gene expression in leaf tissues, presumably reflecting stabilizing selection on gene expression.

## Results

### Differential expression between sexes and life stages

We generated leaf DNA and RNA sequences and pollen RNA sequences from 78 male-female sibling pairs in a single large population (see Methods). We used male and female leaves to compare the genetic architecture of gene expression between the sexes and used male leaves and pollen to compare diploid and haploid life stages, respectively. We performed differential gene expression analyses to identify sex-biased genes in leaves and life stage-biased genes between pollen and male leaf tissues (Table S2). Among the 37,659 annotated genes across the genome, we first filtered for genes with evidence of expression in at least one of the two tissues being compared, resulting in 19,494 genes in the between-sex comparison and 21,337 genes in the between-life-stage comparison. In a principal component analysis (PCA) of RNA read counts, pollen and male leaf samples were separated by PC1 that explained 99% variance both before and after excluding sex-specific regions on sex chromosomes (Fig S1). Male and female leaf samples were separated by PC1 which explained 52% of the variance when sex-specific regions were included, but were not clearly separated based on autosomal and pseudoautosomal (PAR) genes (Fig S1). We identified significantly differentially expressed (DE) genes based on fold change (FC, the ratio of expression level between two tissues) and the false discovery rate corrected *p*-value (adjusted *p*), unless stated otherwise, the cutoffs for DE genes are |log_2_FC| > 1 and adjusted *p* < 0.05. Consistent with the PCA results, there were over 10-fold more DE genes between life stages (16,649) than between sexes (1,496) (Table S2).

We tested for enrichment of DE genes on sex chromosomes and the overlap between sex-biased and life stage-biased genes. Sex-biased genes were highly enriched on sex chromosomes (Table S2), with female-biased genes enriched on the X-specific regions (378/405; Chi-squared test, 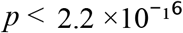) and male-biased genes enriched on the Y-specific regions (1,026/1,091; Chi-squared test, 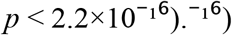 Among 6,702 pollen-biased genes, there was an excess of over-expression of these genes in male leaves compared to female leaves (30 on autosomes and PAR, 1 on X, 269 on Y) relative to female-biased genes (7 on autosomes and PAR, 63 on X, 3 on Y) (Fisher’s exact test, *p* = 0.0028). Although it was a relatively small subset of pollen-based genes, the larger overlap between male leaf-biased and pollen-biased genes was mostly driven by the Y-specific regions which should only be found in males. This finding implies that selection for higher gametophytic expression in pollen could have pleiotropic effects on expression in male leaves (and *vice versa*), which could lead to antagonistic or synergistic pleiotropy for gene expression depending on the direction of selection on expression between life stages.

A previous study in two *Rumex* species found an enrichment of high pollen expression relative to leaves on sex chromosomes compared to autosomes, suggesting a role of male gametophytic selection in Y chromosome evolution (Sandler et al. 2018). With a new PacBio genome assembly containing X and Y phased assemblies (Sacchi et al. 2025), we attempted to replicate these results with our gene expression data in *R. hastatulus*. In accord with the earlier results, we found a significant enrichment of pollen overexpression relative to leaf expression on the Y chromosome compared to autosomes or the X chromosome (Fig S2, Fisher’s exact test, *p* = 0.007 for autosomes, *p* = 0.001 for X) specifically for highly pollen-biased genes (|log_2_FC| > 2, adjusted *p* < 0.05). Among moderately pollen-biased genes (|log_2_FC| > 1, adjusted *p* < 0.05), we found a significant enrichment of pollen-biased genes on the Y chromosome only when compared to the X chromosome (Fig S2, Fisher’s exact test, *p* = 0.031 for X, *p* > 0.1 for autosomes).

### eQTLs between sexes and life stages

To test whether *cis*-eQTLs affect expression concordantly between sexes or life stages, we performed *cis*-eQTL mapping separately in male leaves, female leaves and pollen. For each gene, we tested the nearby SNPs within 20 kb to the transcription start site for significant associations between their genotypes and the focal gene’s expression level, significant associations of gene-SNP pairs after false discovery rate corrections would be considered as *cis*-eQTLs (see Methods). For leaf samples, we mapped *cis*-eQTLs using leaf expression and leaf genotypes in males and females separately. For pollen samples, we used pollen expression and male leaf genotypes as they were from the same male individual and shared the same genotypes. We obtained 4,509,904 and 3,288,995 SNPs on autosomes from 74 female and 69 male leaf DNA samples, respectively, after removing variants with low minor allele frequency or strong Hardy–Weinberg deviations. Based on Tracy-Widom tests, we found 5 and 6 significant PCs for female and male samples explaining 6.53% and 7.7% of genetic variation, respectively (TW statistic ≥ 1.068, *p* ≤ 0.0441 for females; TW statistic ≥ 2.288, *p* ≤ 0.00634 for males); we included the significant PCs as covariates in eQTL mapping.

We identified *cis*-eQTLs and genes whose expression was influenced by eQTLs (“eGenes”) in male leaves, female leaves and pollen (Table 1, Fig S3). We found more shared eGenes and eQTLs between male and female leaves than between male leaves and pollen (Fig 1a-b). The number of eQTLs per eGene ranged from 1 to 126 in pollen, 1 to 170 in male leaves, and 1 to 268 in female leaves, with a median of 2 eQTLs per eGene in both male leaves and pollen, and 3 in female leaves. We found eQTLs were located closer to their associated eGenes than the SNPs tested in all tissues (median distance to the transcription start site: 7718 - 7893 bp for eQTLs vs. 8982 - 9254 bp for tested SNPs). There were similar numbers of eQTLs upstream and downstream of the transcription start site (Fig S4).

**Table 1.** eQTL mapping in male leaves, female leaves and pollen on autosomes. FDR = 0.1.

|  | Number of samples | Number of genes tested | Number of nearby SNPs tested | Number of eGenes | Number of eQTLs | Number of eQTLs affecting multiple genes | Number of secondary eQTLs |
| --- | --- | --- | --- | --- | --- | --- | --- |
| Male leaf | 69 | 14,777 | 1,272,993 | 2,421 | 16,674 | 486 | 367 |
| Female leaf | 74 | 14,564 | 1,687,298 | 3,442 | 34,238 | 905 | 722 |
| Pollen | 69 | 13,785 | 1,182,102 | 1,475 | 9,456 | 201 | 159 |

**Fig 1.**
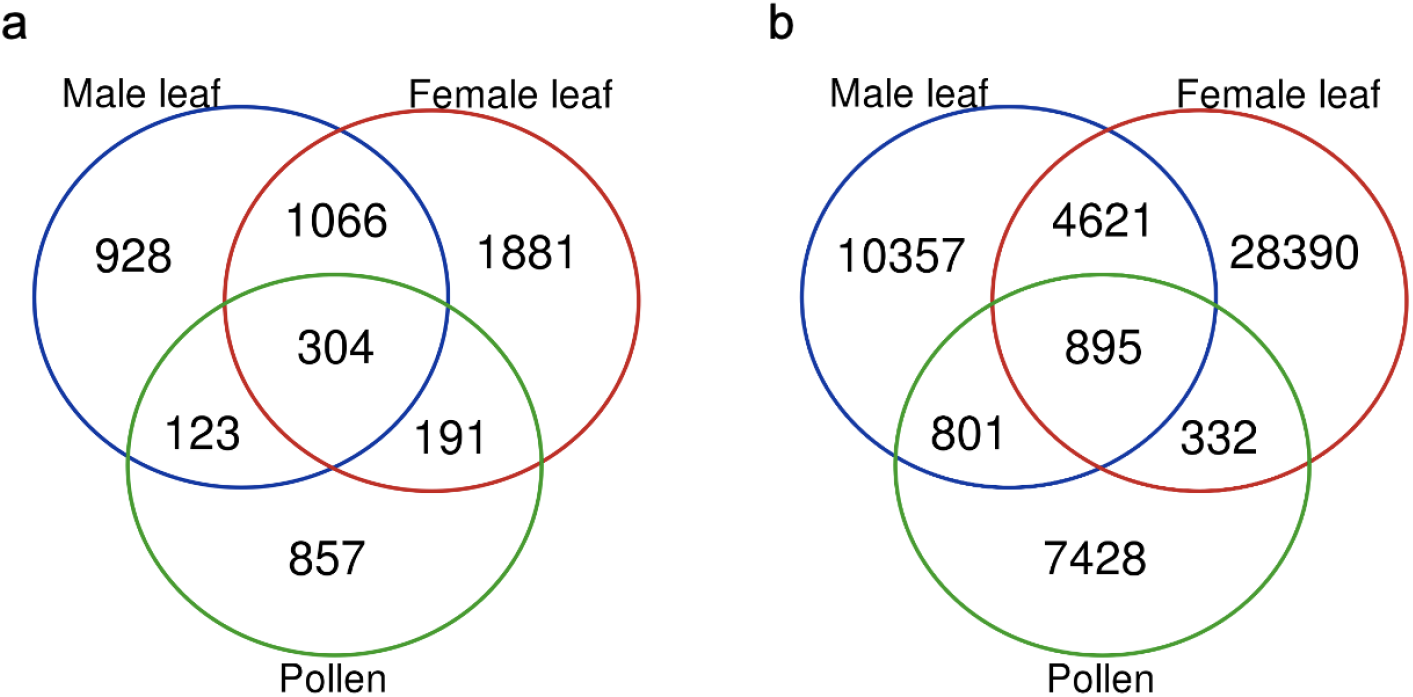
Venn diagrams of eGenes (a) and eQTLs (b) between sexes and life stages on autosomes of *Rumex hastatulus*. FDR = 0.1.

*Cis*-eQTLs with larger effects tend to be closer to the transcription start site (GTEx Consortium et al. 2017). We found a weak but significant negative correlation between effect sizes of eQTLs and their distance to the transcription start site in all tissues when selecting the top eQTL (Spearman correlation, *ρ* = -0.041, -0.0.049, -0.091 for male leaves, female leaves, and pollen, respectively; *p* < 0.0412) and when selecting a random eQTL (*ρ* = -0.067, -0.056, -0.113 for male leaves, female leaves, and pollen, respectively; *p* < 0.001). In addition to the primary eQTLs (usually the top eQTLs) in each tissue, for each eGene we examined the independence of its eQTLs and identified secondary eQTLs (Table 1, see Methods). Secondary eQTLs were considered as conditionally independent after accounting for the effect of the primary eQTL of each eGene. We found primary eQTLs were significantly closer to the transcription start site than secondary eQTLs in female leaf and male leaf tissue (*t*-test, *p* = 0.02, 0.009, 0.32 for female leaves, male leaves, and pollen, respectively). There were 40 - 43% of secondary eQTLs that had opposite effects compared to the primary eQTL of the eGene, consisting of 0.7 - 0.9 % of all eQTLs in each tissue (158 for male leaves, 291 for female leaves, 63 for pollen). The minor allele frequencies of primary and secondary eQTLs were not significantly different (Mann-Whitney *U*-test, *p* = 0.12, 0.37, 0.22 for male leaves, female leaves, and pollen, respectively).

We performed gene ontology enrichment of eGenes in each tissue and examined whether they showed transcriptional or protein biosynthesis functions. Different tissues showed different patterns of functional enrichment (Table S3). For example, female leaf tissue showed enrichment for protein metabolic process (GO:0019538) whereas pollen showed enrichment for mRNA metabolic process (GO:0016071) and negative regulation of gene expression (GO:0010629). Additionally, protein deubiquitination (GO:0016579) was enriched in both male leaves and pollen, and regulation of RNA metabolic process (GO:0051252) was enriched in female leaves and pollen.

### Shared genetic architecture between sexes and life stages

To investigate the extent of shared genetic architecture in expression between sexes and life stages, we first estimated the correlation in expression level across genes between sexes and life stages. For comparing sexes, we estimated the correlation in expression between male and female siblings, which is a phenotypic approximation of the genetic correlation between sexes (*r*_*mf*_). For comparing life stages, this represents the phenotypic correlation between traits, which reflects both environmental and genetic variances. We found expression levels were more positively correlated between sexes than life stages based on all genes and eGenes (Fig 2a-b, Mann-Whitney *U*-test, *p* < 10^-16^ in both comparisons). Even though correlations between life stages involve the same individual (as opposed to siblings of the opposite sex, for comparing the sexes), the stronger correlation in expression between sexes, suggests potentially more shared genetic architecture between sexes than life stages.

**Fig 2.**
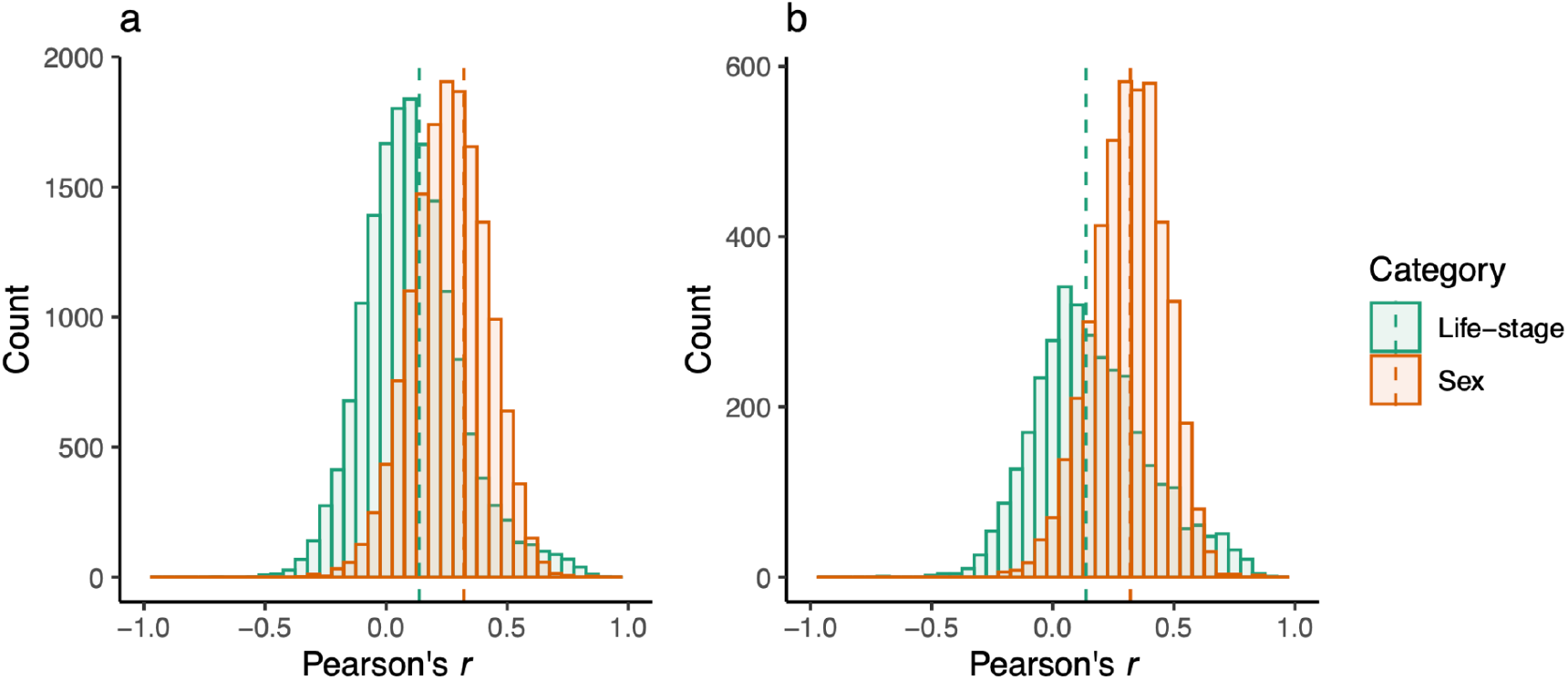
Distribution of Pearson correlation coefficients in gene expression between sexes or life stages in all genes (a) and in eGenes (b) in *Rumex hastatulus*. The dashed line represents the median of each group: 0.093 (Sex) 0.27 (Life-stage) in a, and 0.14 (Sex) 0.32 (Life-stage) in b. Numbers of genes: 14992 (Sex) and 16411 (Life-stage) in a, 4493 (Sex) and 3469 (Life-stage) in b.

Next, we tested whether *cis*-regulatory variation had concordant or discordant effects in gene expression between sexes and life stages. We filtered for nearby SNPs tested in both sexes or life stages that were the top eQTL for the eGene in at least one sex or life stage, resulting in 2,420 eQTLs in the sex comparison and 3,077 eQTLs in the life-stage comparison. The majority of eQTLs showed concordant effects between both sexes and life stages, with their effect sizes positively correlated (Fig 3a-b). The correlation of eQTL effect sizes was much stronger between sexes (Spearman correlation, *ρ* = 0.72, *p* < 10^-16^) than between life stages (*ρ* = 0.35, *p* < 10^-16^), suggesting a stronger shared genetic architecture between sexes (Fig 3a-b), as would be predicted from the phenotypic correlations in Fig 2a-b. There were 170 eQTLs with opposite effects between sexes, all of which were the top eQTL in one sex but not an eQTL in the other sex (light orange and light purple dots in Fig 3a). There were 974 eQTLs with opposite effects between life stages, 19 of which were eQTLs in both life stages (Fig S6). Functional enrichment of eGenes, with eQTLs showing opposite effects between sexes or life stages, showed an enrichment in processes relevant to transcription including negative regulation of gene expression (GO:0010629) and regulation of DNA-templated transcription (GO:0006355), and processes post transcription including regulation of RNA and protein metabolic processes (GO:0051252, GO:0051246) (Table S4).

**Fig 3.**
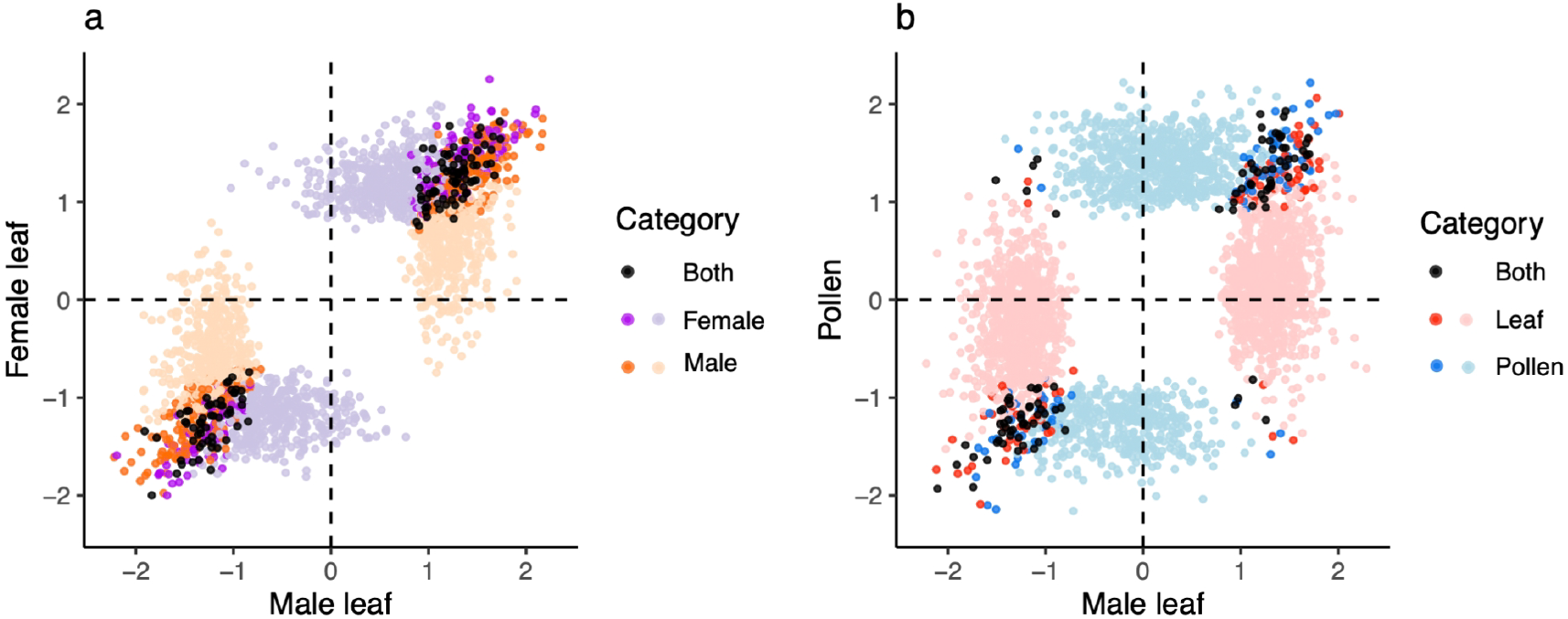
Correlation of effect sizes of autosomal SNPs tested between (a) sexes and (b) life stages in *Rumex hastatulus*. The x and y axes show eQTL effect sizes in different tissues. Black: top eQTL of the eGene in both sexes or life stages; darker colors: top eQTL in the labelled sex/life stage, significant but not the top eQTL in the other sex/life stage; lighter colors: top eQTL in the labelled sex/life stage but not significant in the other sex/life stage. Spearman correlation coefficients *ρ*: 0.72 (a), 0.35 (b).

In addition to mapping *cis*-eQTLs in different tissues separately, we tested for Genotype × Sex interaction effects on expression more explicitly in a linear model using both male and female leaf tissues. We obtained 3,684,937 SNPs on autosomes from 143 leaf DNA samples. We found one significant PC in all leaf samples combined that explained 2% of genetic variation (TW statistic = 1.328, *p* = 0.0302) and we included it as a covariate in eQTL mapping. We tested 14,587 genes and 1,083,706 nearby SNPs for the effect of Genotype, Sex, and Genotype × Sex interaction on expression. After correction for multiple testing (Davis et al. 2016), we found 5 eQTLs showing significant Genotype *×* Sex interaction (adjusted *p* < 0.1, see Methods), all of which changed expression of their eGene in the same direction but with different magnitudes (Fig S7). The small number of eQTLs showing Genotype *×* Sex is consistent with the previous result that eQTLs showed strongly correlated effects between sexes (Fig 3a).

We mapped *cis*-eQTLs affecting the degree of expression differences between life stages using 16,218 genes and 1,404,801 nearby SNPs. Specifically, we aimed to test the effects of the minor allele of each SNP on the differences in expression levels between male leaf tissue and pollen. We found 9,873 eQTLs affecting the expression differences between life stages in 1,564 genes. There were more eQTLs in which the minor allele increased expression differences (6,173) than reduced them (3,700), suggesting new mutations (typically represented by the minor alleles) more often drive the divergence in expression between life stages than constrain it. We examined whether eQTLs for expression differences between life stages were also identified as eQTLs in male leaf tissue or pollen separately. We found 767 eQTLs significantly affected leaf expression, pollen expression, and the differences between them, 735 of which affected leaf and pollen expression in the same direction, consistent with the concordance in eQTL effect sizes shown in Fig 3b. Additionally, we found that 1,464 of 1,564 genes, whose expression differences between life stages were affected by eQTLs, were significantly DE genes between pollen and male leaf tissue (adjust *p* < 0.05, no FC cutoff). Functional analyses showed rRNA processing (GO:0006364) and regulation of RNA metabolic processes (GO:0051252) were enriched in genes with eQTLs affecting life-stage expression differences (Table S5). Lastly, we identified 141 conditionally independent eQTLs affecting life-stage expression differences, 54 of which had opposite effects compared to the primary eQTL of the same gene.

### Selective pressure on eQTLs

We examined the distribution of minor allele frequencies (MAF) of eQTLs to test if selective pressures differed for eQTLs between sexes or life stages. We first tested if eQTLs show more common or rare alleles, and whether the distributions of MAFs differed across tissues. We kept one eQTL per eGene to have a set of independent eQTLs for MAF comparisons and repeated analyses using both the top eQTL and a randomly selected eQTL. In all tissues, there were higher proportions of rare alleles than intermediate-frequency alleles regardless of the eQTL selection method (Fig 4a-f). The median MAFs were similar across tissues, with the highest in female leaves and lowest in male leaves using both eQTL selection methods (Fig 4). The differences in mean MAFs between tissues were marginally significant between male and female leaves (Mann-Whitney *U*-test, top eQTL: *p* = 0.041, random eQTL: *p* = 0.049) and between female leaves and pollen in top eQTLs (*p* = 0.049) (Fig 4).

**Fig 4.**
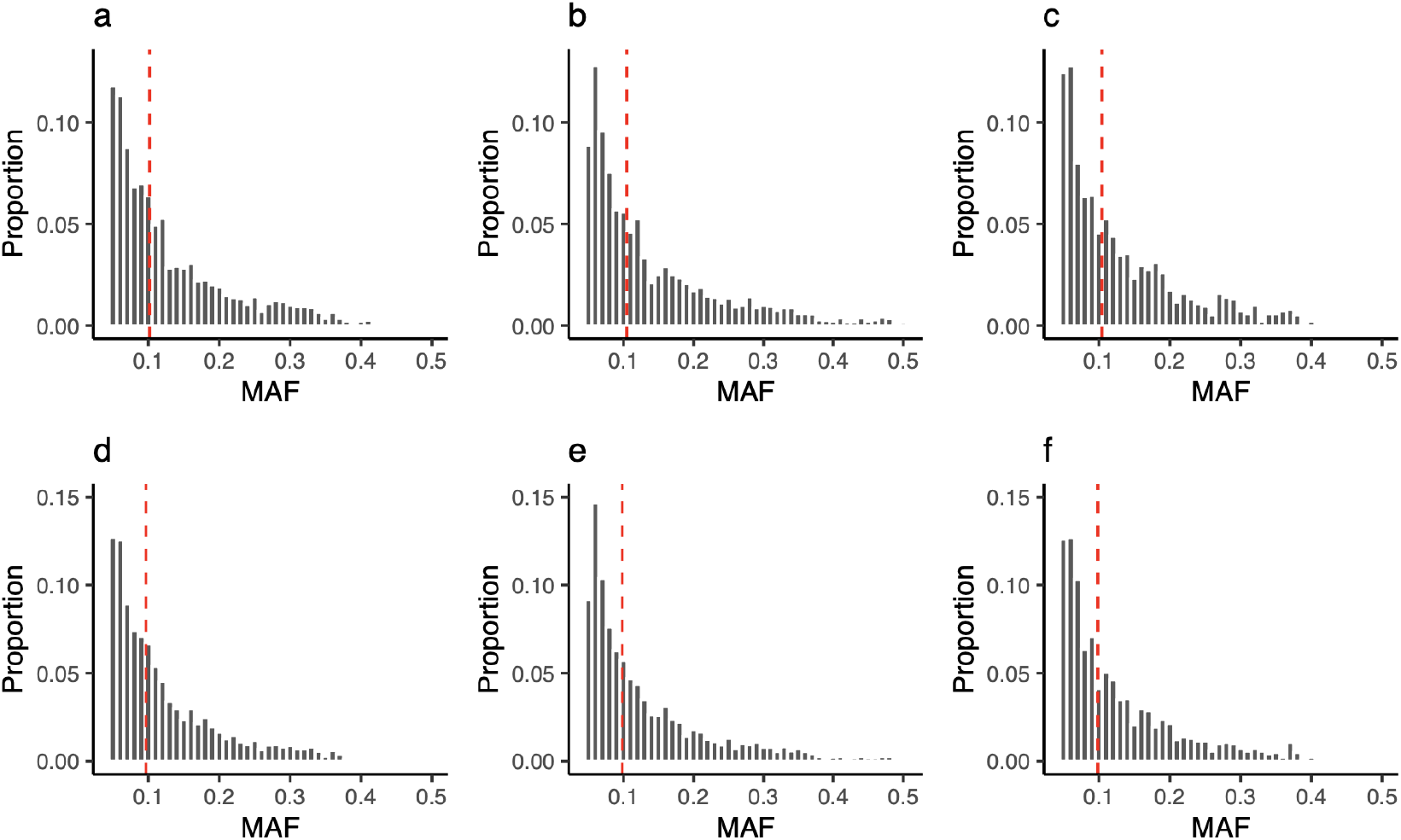
Distribution of MAFs of eQTLs in male leaves (a, d), female leaves (b, e), pollen (c, f) in *Rumex hastatulus*. a-c: top eQTL per eGene, d-e: a randomly selected eQTL per eGene. Binwidth = 0.01. Dashed lines represent the Median MAF in each plot: 0.1017 (a), 0.1048 (b), 0.1045 (c), 0.0965 (d), 0.0984 (e), 0.0982 (f).

Under purifying selection, new mutations affecting gene expression would be predominantly deleterious and removed from the population resulting in an excess of rare alleles compared to neutral expectations. To investigate whether the ‘excess’ of rare alleles in eQTLs across tissues in Fig 4 reflects purifying selection relative to a neutral standard, we compared the MAFs of true-positive eQTLs to a null distribution of false-positive eQTLs from permutated data, generated by shuffling the assignment of expression phenotypes to individuals (see Methods).

We identified false-positive eQTLs from permuted data using the same methods as the observed data. In all tissues, the permuted data showed a higher proportion of rare alleles (MAF 0.05 - 0.1) than the observed data, with only a small overlap in pollen (Fig 5a-c). This result suggests that eQTLs were not enriched for rare alleles relative to the null distribution and thus does not support a genome-wide signal of purifying selection on *cis*-regulatory variation in *R. hastatulus*. We also compared MAFs of eQTLs shared between life stages and those specific to one life stage, and MAFs of eQTLs on the X-specific and pseudo-autosomal regions in female leaf (see Supplementary Results).

**Fig 5.**
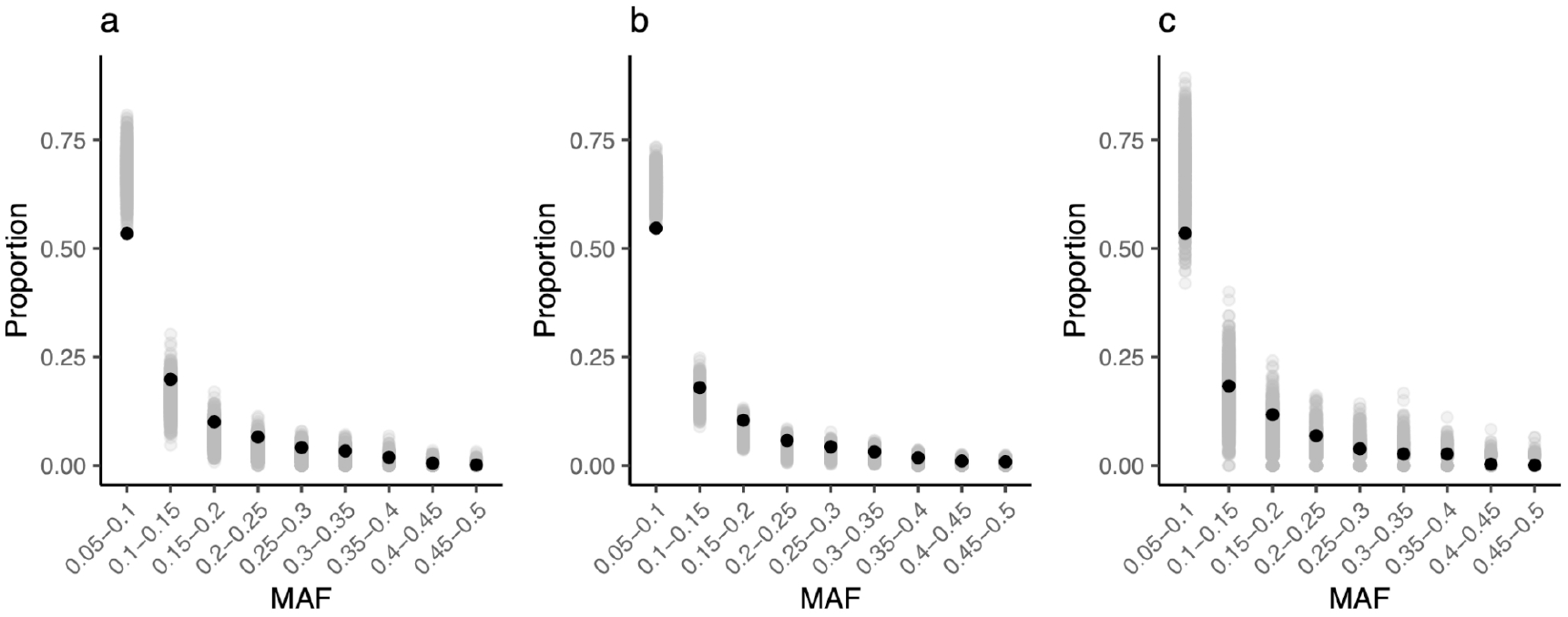
Distribution of MAFs of eQTLs compared to a null distribution in male leaves (a), female leaves (b) and pollen (c) in *Rumex hastatulus*. Black dots: true positive eQTLs (observed data), grey dots: false positive eQTLs (permuted data).

Under antagonistic selection, genes shared between sexes or life stages should have higher genetic diversity due to a greater potential for ongoing conflict compared to sex-specific or life-stage-specific genes. To test for antagonistic selection in coding regions, we compared neutral diversity statistics such as *π* and Tajima’s in different groups of eGenes using 4-fold degenerate sites which are often assumed to be neutral. We predicted that shared eGenes would have higher *π* and Tajima’s *D* than specific eGenes. We found that shared eGenes between sexes had significantly higher Tajima’s *D* (Kruskal-Wallis test, *p* < 10^-5^) but not significantly higher *π* (*p* = 0.5383) than male- or female-specific eGenes. However, the elevated Tajima’s *D* may be driven by ascertainment bias where higher allele frequencies increase the power to detect eQTLs, thus shared eGenes have higher diversity regardless of selection. The ascertainment bias makes it difficult to conclude that shared eGenes are more likely to undergo antagonistic selection. We found no significant differences in *π* or Tajima’s *D* between shared and specific eGenes in either life stage (Kruskal-Wallis test, *p* = 0.8586, 0.2374, respectively). Additionally, eGenes whose expression differences between life stages were affected by eQTLs had significantly higher *π* and Tajima’s *D* than other eGenes (*t*-test, *p* = 0.00013, 0.00058, respectively).

### Burden of rare alleles on gene expression

While our comparisons of allele frequencies of eQTLs to the null distribution suggest a deficiency of rare variants rather than a signal of purifying selection on variants affecting gene expression, the rarest variants are excluded in eQTL mapping due to insufficient power to detect associations. To address this limitation, we performed burden of rare allele tests that assess signatures of stabilizing selection in gene expression (Zhao et al. 2016; Kremling et al. 2018; Uzunović et al. 2019). This test evaluates the shape of the distribution of total counts of rare alleles across ranked bins of varying expression levels. Under stabilizing selection against extreme expression, individuals at highest or lowest expression ranks will carry more rare alleles, driving a parabolic pattern. In *Capsella grandiflora* leaf tissues, rare SNPs are equally likely to increase and decrease expression (Uzunović et al. 2019); whereas in maize tissues including leaves, roots, shoots and kernels, rare SNPs upstream of genes are more likely to decrease expression (Kremling et al. 2018). We ranked individuals’ expression levels for each gene and counted the number of singletons (i.e., variants that only appear once across all individuals) in the 5kb upstream window of the gene across all genes for each expression rank to evaluate the patterns in *R. hastatulus* (Fig 6).

**Fig 6.**
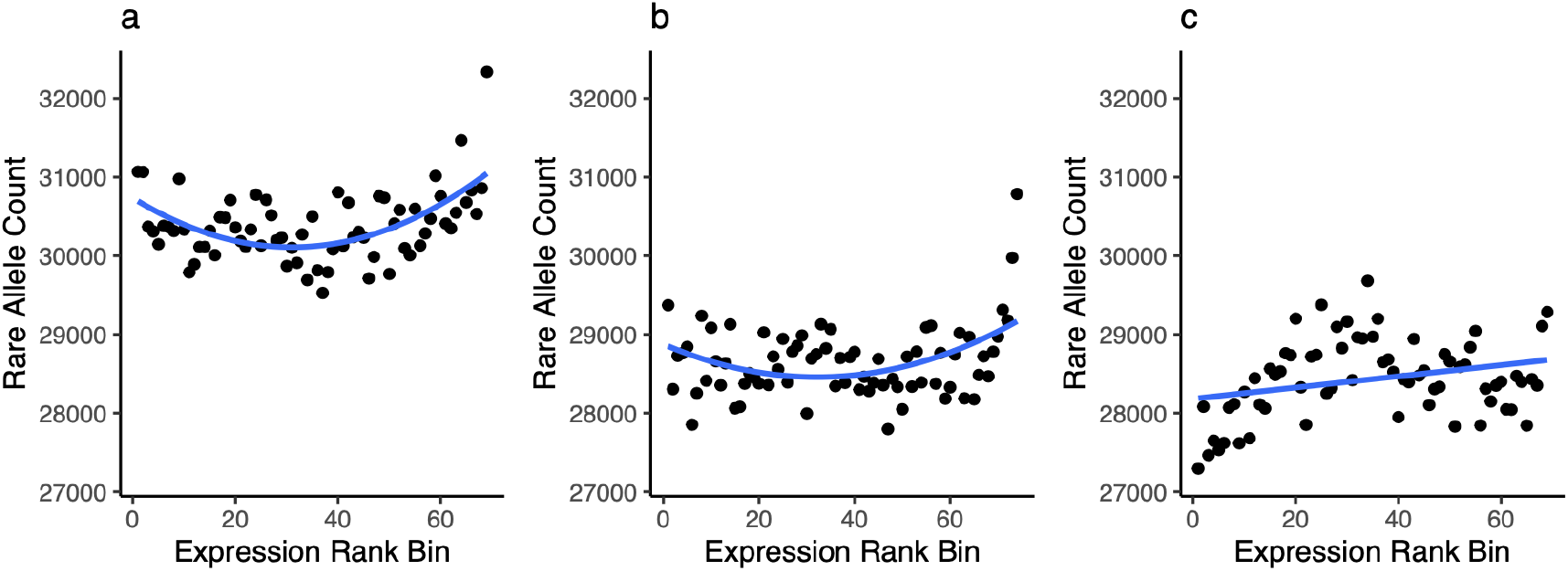
Relationship between rare allele count and gene expression rank bin in male leaves (a), female leaves (b) and pollen (c) in *Rumex hastatulus*. Smooth lines show quadratic regression in a and b and linear regression in c. a. adjusted *R*^2^ = 0.5288, *p* = 6.149×10^-12^. b. adjusted *R*^2^ = 0.5484, *p* = 2.067×10^-13^. c. adjusted *R*^2^ = 0.3481, *p* = 5.756×10^-8^. Lowest rank means lowest expression and highest rank means highest expression.

In male and female leaves, rare alleles were associated with both higher and lower gene expression (Fig 6a-b). The relationship between rare allele count and expression rank in leaves appeared to be parabolic and overall symmetrical, with only a few outliers in the highest expression ranks (Fig 6a-b). This parabolic pattern is consistent with stabilizing selection favoring intermediate levels of expression, similar to the results in *Capsella grandiflora* leaf tissues (Uzunović et al. 2019). In pollen, we did not find the same parabolic pattern (Fig 6c); both quadratic and linear regressions were significant, but the linear model (adjusted *R*^2^ = 0.3481, *p* = 5.756×10^-8^) explained more variance compared to the quadratic model (adjusted *R*^2^ = 0.2979, *p* = 3.181×10^-6^). The lack of a parabolic pattern provides little evidence for stabilizing selection in pollen, in contrast with leaves.

## Discussion

In this study we mapped *cis*-eQTLs in leaf and pollen tissues in the dioecious plant *R. hastatulus* to investigate the potential for conflict between sexes and life stages, based on shared genetic architecture. We investigated the overlap of eQTLs between sexes and life stages and examined their effect sizes and allele frequencies. We discovered a much larger overlap in eGenes and eQTLs between sexes than between life stages. For SNPs tested in both sexes or life stages, there was a stronger correlation of effect sizes between sexes suggesting a more shared genetic architecture between sexes than life stages. There were more minor alleles of eQTLs driving than constraining divergent expression between life stages. Among the eQTLs shared between life stages and affecting their expression differences, most minor alleles changed expression in the same direction with different magnitudes, increasing the degree of divergent expression. MAFs of eQTLs in all tissues exhibited similar distributions with a larger proportion of rare alleles than common alleles, but this pattern did not reflect a clear genome-wide signal of purifying selection when compared to a null distribution. Investigation of the burden of rare allele on genotypes with more extreme gene expression suggested that rare variants both up- and down-regulated expression in leaves but not in pollen. This finding is consistent with stabilizing selection on gene expression in leaves only. Below we discuss the implications of our results on the extent of a shared genetic architecture between sex and life stages and also the use of eQTL mapping in testing evolutionary forces shaping *cis*-regulatory variation.

### Potential for conflict between sexes and life stages

With the potential for both sexual and life-stage conflict based on theory (Kidwell et al. 1977; Immler et al. 2012; Peters and Weis 2018), key open questions remain on whether there are more opportunities for conflict between sexes or life stages, and whether sexual and life-stage conflict is still an ongoing process or has already become resolved. We investigated these questions using *cis*-regulatory variation which has a strong effect on gene expression. Our results consistently suggested significantly greater shared genetic architecture between sexes than life stages, thus more potential for ongoing sexual conflict. Furthermore, our results indicated that life-stage conflict is more resolved compared to sexual conflict. These findings are consistent with previous work suggesting considerable scope for ongoing sexual conflict based on intersexual genetic covariances in expression (Allen et al. 2018 2018; Houle and Cheng 2021; Singh and Agrawal 2023; Grieshop et al. 2025).

It is also worth mentioning that standing variation in gene regulation might have more sex-differential effects in reproductive tissues such as in flowers compared to leaves. Future studies could benefit from a more diverse sampling of both vegetative and reproductive tissues (e.g., comparing pollen and ovule or shared floral tissues) than we used in this study. The alternating life cycle of diploid and haploid life stages is shared among all land plants, allowing adaptation to varying environments. While sexual reproduction is an ancient attribute of eukaryotic organisms, dioecy in flowering plants has more recently emerged from hermaphroditic ancestors in most lineages (Bawa 1980), and there probably has not been enough time to break down intersexual genetic correlations. Our conclusion that there is more potential for sexual than life-stage conflict in dioecious flowering plants should be robust in terms of tissue sampling due to the relatively recent evolution of separate sexes compared to the fundamental feature of the haploid-diploid alternating life cycle. In dioecious plants such as *R. hastatulus*, it is likely that there has been a greater amount of time to resolve conflicts between life stages than sexes, where sexual conflict between females and males may still be ongoing. Direct assessments of antagonistic selection between sexes and life stages should provide a more complete picture of the evolution of these two types of conflict.

While our results suggest the potential for unresolved sexual conflict in gene expression on autosomes, sex chromosomes have important roles in the evolution of dioecy (Charlesworth et al. 2005) and can facilitate the evolution of sexual dimorphism (Rice 1984; Zemp et al. 2016). The vast majority of identified sex-biased genes were found to be sex-linked, suggesting a potential role for sex chromosomes in resolving sexual antagonism. Plants are useful systems for comparing opportunities for sexual and life-stage conflict (Moore and Pannell 2011; Beaudry et al. 2020). Whether there is more shared genetic architecture between sexes and life stages in plant lineages with different reproductive systems or sex chromosome systems (presence and absence or age of sex chromosomes) is therefore worth further investigation. Also, whether there are differences in sexual and life-stage conflict in plants with contrasting sexual systems (hermaphroditic vs. dioecious) or different types of life cycles (predominantly diploid vs. haploid) are also promising topics for future research.

With a weak positive correlation of eQTL effect sizes between life stages there is still potential, albeit weaker, for ongoing life-stage conflict. We found that genes on the Y chromosome with higher expression in pollen compared to leaves also tended to be male-biased in leaves, suggesting the possibility of strong selection in pollen, possibly driving up leaf expression in an antagonistic fashion. We examined signatures of conflict on eGenes using diversity statistics. We expected eQTLs with concordant effects between sexes or life stages to experience stronger conflict than those with discordant effects or those exclusive to one sex or life stage. We found that shared eGenes had higher Tajima’s *D* than sex-specific eGenes. In contrast to our prediction, eGenes with eQTLs affecting their expression differences between life stages had significantly higher *π* and Tajima’s *D* than eGenes not significant for expression differences; however, this finding might result from an ascertainment bias, as it is more likely to have higher power to detect interactions when diversity is higher. Overall, there was no indication of greater antagonistic selection on eQTLs with shared genetic architecture between the sexes and life stages.

Our previous study tested the genomic signature of intra-locus conflict between life stages in *R. hastatulus* using gene expression but did not find support for a genome-wide signal (Yuan et al. 2025). Conditions for generating balancing selection due to conflict are limited to single large-effect loci (Kidwell et al. 1977). Detecting balancing selection due to conflict using genomic data remains challenging, especially when polygenic traits are involved (Bitarello et al. 2023; Flintham et al. 2025; Ruzicka et al. 2025). While variation in gene expression can underlie important phenotypic variation, it is worth noting that the relationship between expression variation and morphological variation can be complicated and different across tissues. Clearly, there is more to be understood about the relationship among gene expression, phenotypic traits and fitness. For example, the genetic basis of the traits under antagonistic selection needs to be considered when testing for signals of conflict (Rowe et al. 2018). Future studies with more functional information (e.g., on alleles beneficial for pollen competition), combined with previously identified candidate genes under balancing selection (Yuan et al. 2025) should help to uncover the genetic basis of life-stage conflict in *R. hastatulus*.

### Selective pressure on cis-regulatory variation

Estimating selective pressures in regulatory variation using allele frequencies has resulted in mixed results supporting contrasting hypotheses about the type of selection on *cis*-eQTLs (Josephs et al. 2015; Brown and Kelly 2021). Our results indicated more rare than common alleles among *cis*-eQTLs in all tissues. However, the proportion of rare alleles in the observed data was lower than the null distribution generated from the permuted data, thus we found no evidence for a genome-wide signal of purifying selection. Our results are intermediate between those reported in *Capsella grandiflora* supporting purifying selection (Josephs et al. 2015) and those in *Mimulus* floral tissues, which showed a higher proportion of common than rare alleles (Brown and Kelly 2021). The slight deficiency of rare alleles relative to the null distribution could potentially be caused by antagonistic balancing selection maintaining variation at intermediate frequencies or weaker purifying selection on detectable eQTLs. Our current data cannot distinguish these alternative explanations, partly because the genome-wide null distribution likely includes a sampling of variants subject to purifying selection and does not reflect a strictly neutral baseline for comparisons.

The burden of rare allele test provides a complimentary and more powerful assessment of selective pressure on genetic variation affecting gene expression, as it captures effects of the rarest variants excluded from eQTL mapping studies due to low statistical power. Using this approach, we found a clear signal of stabilizing selection on gene expression in male and female leaves, with an excess of rare allele burdens at both extremes of expression rank, suggesting selection against extreme expression as purifying selection keeps rare deleterious alleles at low frequencies at extreme expression levels (Hodgins-Davis et al. 2015; Uzunović et al. 2019). In contrast, we observed no signal of stabilizing selection on gene expression in pollen. This may indicate that pollen expression is generally subject to weaker selection in *R. hastatulus*, a conclusion that is supported by our recent results suggesting relaxed purifying selection on pollen-biased genes (Yuan et al. 2025). Future studies should benefit from examining the correlation between effect sizes and MAFs, while controlling for confirmation bias using allele-specific eQTLs (Josephs et al. 2015) and examining *trans*-eQTLs as they have been found in other species to be subject to stronger constraint than *cis*-acting variation (Josephs et al. 2020).

In conclusion, our results indicate a greater shared genetic architecture in gene expression between the sexes than between the haploid and diploid phases of the lifecycle, and evidence for stabilizing selection on gene expression in leaves but not pollen. These results imply more potential for ongoing conflict between sexes than life stages in *R. hastatulus*, though the extent of standing sexually antagonistic variation remains difficult to quantify from our current data alone. Future work using more direct mapping of fitness effects of genetic variants (e.g., Ruzicka et al. 2019; Delph et al. 2022) will be important to test this further.

## Methods

### Plant materials

We described the details of plant materials and sequencing in Yuan et al. (2025); here we provide a brief summary and additional relevant information. After generating seedlings using open-pollinated seeds collected from a population in Rosebud, Texas, US (Pickup and Barrett 2013), we paired one female and male individual randomly at the time of flowering for crossing and collected seeds to generate F_1_ plants. We planted and retained multiple individuals per family for 78 F_1_ families to ensure we had one male-female full sibling pair per family for tissue collection. We collected young leaf tissue from each male-female full-sibling pair after flowering for both RNA and DNA sequencing and collected mature pollen grains from males for RNA sequencing. We described the DNA and RNA isolation protocols in Yuan et al. (2025). The DNA samples were sequenced at a depth of 10-15X. Raw sequences of male leaf RNA and pollen RNA were described in Yuan et al. (2025), Initial sample sizes for DNA samples and female leaf RNA samples are presented in Table S1.

### DNA Seq data and variant calling

We used a phased PacBio genome assembly consisting of four autosomes, X-specific, Y-specific and pseudoautosomal regions for our analyses (Sacchi et al. 2025). We mapped the DNA reads to the genome assembly and added read groups using bwa-mem2 (Vasimuddin et al. 2019). We sorted the BAM, marked PCR duplicates and indexed the final BAM using SAMTools (Danecek et al. 2021). We called variants in parallel on 2 Mb non-overlapping contigs of each chromosome for all samples using BCFtools mpileup (Danecek et al. 2021), then concatenated the VCFs of contigs for each chromosome. We removed one male sample (24fM) due to low genomic coverage and two samples due to incorrect sex labeling (35aM, 40aF), and only kept samples with both DNA and RNA sequences from the same individual. For males, we only kept samples with both leaf and pollen RNA sequences from the same individual. Our sample sizes for VCF filtering were 74 for female leaves, 73 for male leaves, making a total of 147 leaf tissue samples. We performed SNP filtering as follows for female leaf, male leaf and all leaf samples combined separately using VCFtools (Danecek et al. 2011). We kept SNPs with QUAL > 30, a mean depth between 5 and 20, a genotype quality per sample > 30 otherwise marked as missing, and finally a missingness < 20%. For female samples, we used the same filtering criteria for autosomes, PAR and X-specific regions. The numbers of autosomal SNPs were 64,988,762 in male leaf, 66,828,708 in female leaf, and 66,341,762 in all leaf samples combined. The numbers of SNPs on X-specific regions and PAR in females were 7,745,941 and 2,164,214, respectively. Additionally, we filtered for invariant sites on autosomes with a mean depth between 5 and 20 and a missingness < 20% for all leaf samples combined.

We removed SNPs with a minor allele frequency lower than 5% or with strong Hardy–Weinberg deviations (*p* < 10^-6^) on autosomes using Plink (Chang et al. 2015) to test for population structure and for the subsequent eQTL mapping. We removed four male samples (7bM, 27eM, 53bM, 5aM) from subsequent analyses for being outliers in principal component analyses in Plink (Chang et al. 2015). Our final sample sizes were 74 for female leaves, 69 for male leaves, making a total of 143 leaf samples. After the filtering using Plink, the numbers of SNPs on autosomes were 3,288,995 in male leaves, 4,509,904 in female leaves, and 3,684,937 in all leaf samples combined; the numbers of SNPs on X-specific regions and PAR in females were 769,059 and 96,911, respectively.

### RNA Seq data and expression analyses

We mapped the RNA reads to our genome assembly in a two-pass mode, sorted the BAM files by coordinates, added read groups using STAR (Dobin et al. 2013), and indexed the output BAM files using SAMTools (Danecek et al. 2021). We generated read counts for each gene using featureCount (Liao et al. 2014). We only kept genes with evidence of expression for our subsequent analyses, i.e., genes with a mean raw read count ≥ 5 in a specific tissue, or in at least one tissue for analyses involving both sexes or life stages. We identified sex-biased genes in leaf tissues and life-stage-biased genes by comparing pollen and male leaf tissues using DESeq2 (Love et al. 2014). The cutoffs for significant differentially expressed (DE) genes were Benjamini-Hochberg adjusted *p* < 0.05 and |log_2_ FC| > 1 unless stated otherwise.

We prepared the expression phenotype data for each tissue separately and for all leaf samples combined for eQTL mapping. We normalized raw read counts by sequencing depth using DESeq2 (Love et al. 2014) and performed quantile normalization using the qqnorm function in R (R Core Team 2022). Additionally, we calculated the difference of normalized read counts in pollen and male leaves (male leaf minus pollen) for each gene and performed quantile normalization of its absolute value using the qqnorm function in R (R Core Team 2022).

### Correlation in gene expression

To characterize the correlation in expression across life stages or sexes, we used Pearson correlations and the final normalized read counts as expression level as described above. For male leaves and pollen, we calculated the correlation of a given gene’s expression level in each tissue using 69 male individuals and repeated for 16,411 autosomal genes that had evidence of expression (as described above) to generate a distribution of correlations. These correlations are the same as any phenotypic correlation in behavioral or morphological traits, but here are for gene expression (Stinchcombe and Kelly 2025). To estimate the across-sex correlation in expression, we took advantage of the sib-structure of our design and the fact that we had male and female sibling pairs. We used the expression level of a gene in a male sample and its female sibling as a pair of observations, repeated for 64 sibling pairs in our design, to estimate the correlation in expression across siblings for a given gene. We then repeated this for 14,992 genes with evidence of expression in the samples to generate a distribution of across-sex correlation in expression. Note that because we only had a single male and female sample per family, rather than replicates, this is a phenotypic approximation of the across-sex genetic correlation (*r*_*mf*_), rather than a genetic correlation. We repeated the analyses for genes regulated by eQTLs (eGenes, see below) in at least one life stage or sex following the same steps.

### eQTL mapping

We performed *cis*-eQTL mapping on autosomes separately for male leaves, female leaves, and pollen tissue. For female leaves, we repeated eQTL mapping on X and PAR using the same approach as the autosomes. For male samples, we performed eQTL mapping using the difference in leaf and pollen expression levels as phenotypes in the same way as in leaves or pollen separately. To control for genetic relatedness, we performed LD pruning in Plink (Chang et al. 2015) and identified genetic principal components (PC) using EIGENSOFT smartpca (Patterson et al. 2006; Price et al. 2006). The significance of each PC was determined by the Tracy-Widom test (Patterson et al. 2006; Sztepanacz and Blows 2017). We included the significant PCs (*p* < 0.05) as covariates in eQTL mapping.

We mapped *cis*-eQTL using tensorQTL (Taylor-Weiner et al. 2019) following its guidelines, and tested pre linkage-pruning SNPs within a 20 kb window to the transcription start site for each gene. We performed adaptive permutations to calculate empirical and beta-approximated empirical *p*-values for each gene (Ongen et al. 2016), and generated nominal *p*-values for all SNP-gene pairs. We calculated the Storey *q*-value and the nominal *p*-value threshold for significant associations based on the beta-approximated empirical *p*-value for each gene using a false discovery rate (FDR) of 0.1 (Benjamini and Hochberg 1995; Storey 2002). We filtered for genes that had *q*-value < 0.1 and at least one eQTL as our eGenes. We filtered for SNPs with a nominal *p*-value smaller than the nominal *p*-value threshold of its eGene as our eQTLs. The minimum *p*-value threshold is 0.0318 for male leaves, 0.0503 for female leaves, and 0.0166 for pollen. We identified conditionally independent eQTLs for eGenes using a forward-backward stepwise regression model in tensorQTL (GTEx Consortium et al. 2017). We defined primary eQTLs as the top ranked eQTL for each eGene, and any additionally ranked eQTLs as secondary or conditionally independent eQTLs. The effect size of an eQTL was defined as the slope in the linear regression model from tensorQTL. In the analyses of eQTLs for differences in expression levels between life stages, effects of the minor allele were tested rather than alternative allele of each SNP.

We kept one eQTL for every eGene, either the most significant (top) eQTL or a randomly selected eQTL when comparing minor allele frequencies (MAF) and when testing the relationship between effect size and the distance to transcription start site of eQTLs. We removed eQTLs and eGenes when an eQTL affected multiple eGenes when comparing MAFs. We performed gene ontology enrichment of eGenes using topGO (Alexa and Rahnenfuhrer 2023), significant GO terms were selected using a *p*-value cutoff of 0.05 based on Fisher’s exact tests.

We ran tensorQTL on all 143 leaf samples to test the linear model: Expression ∼ Genotype + Sex + Genotype × Sex. We kept the top association for each gene based on the interaction term (-best only), and calculated multiple-testing corrected (Davis et al. 2016) and Benjamini-Hochberg adjusted *p*-values (pval_adj_bh) following the recommendations from tensorQTL. The cutoff for eQTLs showing significant Genotype × Sex is pval_adj_bh < 0.1.

### Null distributions of eQTL MAFs

To generate a null distribution for MAFs of eQTLs in each tissue, we performed permutations by randomly re-assigning sample ID’s in the expression phenotype file. We performed 1000 permutations in both females and males. For female leaves, we performed the same permutations on X and PAR as the autosomes. For male samples, we performed permutations the same way in male leaf and pollen phenotypes and identified the shared and specific eQTLs between male leaves and pollen in each permutation. We identified false-positive eQTLs using the same *p*-value thresholds from the previous analyses on our observed data. We removed eGenes with more than 10 eQTLs, randomly selected an eQTL from each eGene, and removed eQTLs affecting multiple eGenes. We repeated the same procedure of calculating MAF using true positive eQTLs. The proportions of false positive eGenes compared to true positive eGenes were 0.059 in male leaves, 0.105 in female leaves, and 0.0318 in pollen, based on the median number of false positive eGenes in 1000 permutations.

### Diversity statistics

We used the script codingSiteTypes.py (https://github.com/simonhmartin/genomics_general/blob/master/codingSiteTypes.py, accessed in 2020) to extract 4-fold degenerate site for calculating neutral diversity statistics. We generated 4-fold VCFs including both variant and invariant sites using all leaf samples combined. We used pixy (Korunes and Samuk 2021; Bailey et al. 2025) to calculate nucleotide diversity (*π*) and Tajima’s *D* for all autosomal genes. We only kept genes with at least 50 sites in the subsequent analyses of *π* and Tajima’s *D*. We removed genes if they were associated with an eQTL that affected multiple eGenes.

### Burden of rare allele test

We performed the burden of rare allele test separately for male leaves, female leaves and pollen following Zhao et al. (2016). We counted the number of singletons in the 5kb upstream window from the transcription start site for each gene and each individual. We then ranked the individuals based on their expression level for each gene, we summed the number of local singletons from all genes for each rank regardless of their individual identity. Finally, we plotted the sum of rare alleles against rank bins (bin size = 1) and tested for quadratic regression in all tissues to assess the parabolic relationship between rare allele counts and gene expression ranks. In pollen, where no clear parabolic pattern was observed, we additionally tested for linear regression and compared adjusted *R*^*2*^ from both models to determine which better describes the data.

## Data availability

Raw sequencing reads are available through the NCBI SRA database under BioProject PRJNA744278. Scripts used in this study are available at https://github.com/SIWLab/Rumex_eQTL.

## Acknowledgement

We thank Yunchen Gong for technical assistance with the computing servers and Bill Cole and Thomas Gludovacz for help with plant maintenance. This study was supported by NSERC Discovery Grants awarded to SCHB, JRS and SIW.

