## Supplementary Materials for "*Cis*-regulation of gene expression between sexes and life stages in *Rumex hastatulus*"

### Supplementary Results

*Selective pressures on shared eQTLs between life stages*

Under antagonistic selection between life stages, eQTLs shared between life stages are expected to have higher allele frequencies than life-stage-specific eQTLs, as antagonistic pleiotropy can maintain shared genetic variation at intermediate frequencies. We tested this prediction by comparing MAFs of shared versus life-stage-specific eQTLs. We did not perform the same comparison between sexes because eQTLs in male leaf and pollen were derived from the same individuals and share the same allele frequencies, whereas male and female leaf eQTLs always have different allele frequencies and a similar comparison is therefore not possible. Consistent with our prediction, we found shared eQTLs had significantly higher MAFs than specific eQTLs between life stages (Mann-Whitney $U$-test, $p < 10^{-7}$), with median MAFs of 0.143 and 0.093, respectively. However, we could not confirm this result against a null distribution due to a lack of shared false-positive eQTLs between life stages as the probability of the same SNP being a false positive in both tissues is extremely low. Although the direction of this result is consistent with antagonistic pleiotropy, we cannot rule out the possibility that shared eQTLs have greater power to be mapped in both tissues, thus having higher MAFs. Studies on *Drosophila melanogaster* have found more *cis*-regulatory variation in unbiased and moderately sex-biased genes than in strongly biased genes, suggesting potential sexual conflict in regulatory variation (Puixeu et al. 2023; but see Mishra et al. 2024); whether a similar pattern holds between life stages needs to be further tested.

*eQTLs on sex chromosomes in female leaf*

We identified 111 eGenes with 714 eQTLs in the PAR, and 1,294 eGenes with 16,112 eQTLs in the X-specific regions in female leaf and compared their selective pressure using MAFs. The MAFs of eQTLs in PAR and the X-specific regions showed more rare than common alleles similar to the autosomes (Fig S8). We found eQTLs in the X-specific regions had significantly lower MAFs than in autosomes using both methods of eQTL selection (Mann-Whitney *U*-test, $p < 0.004$), and eQTLs on PAR had significantly higher MAFs than the X-specific regions only in top eQTLs (Mann-Whitney *U*-test, $p = 0.019$). However, the null distribution of MAFs of eQTLs for X and autosomes did not differ significantly (Fig S9); there were not enough false-positive eQTLs in the PAR for comparison.

**Supplementary Tables**

Table S1. DNA and RNA samples of *Rumex hastatulus* used in this study.

Table S2. Number of DE genes in *Rumex hastatulus*.

Table S3. GO enrichment of eGenes in male leaf, female leaf, and pollen of *Rumex hastatulus*.

Table S4. GO enrichment of eGenes with discordant eQTLs between life-stages or sexes in *Rumex hastatulus*.

Table S5. GO enrichment of eGenes with discordant eQTLs for expression differences between life-stages of *Rumex hastatulus*.

**Supplementary Figures**

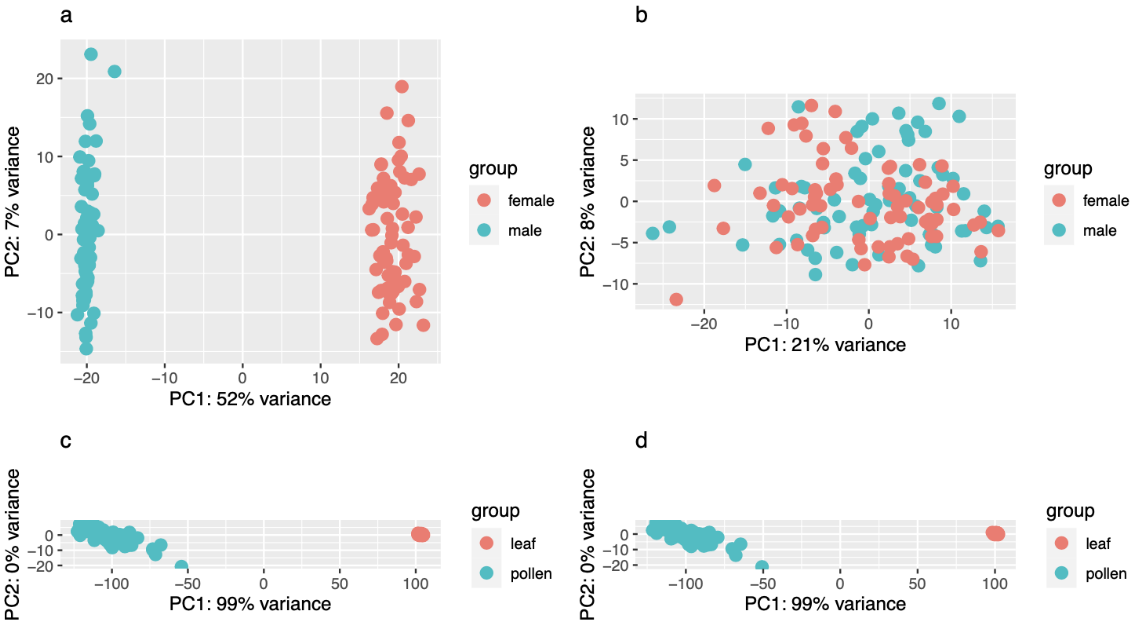


Fig S1. PCA of leaf and pollen RNA samples of *Rumex hastatulus*. a-b: male vs. female leaf. c-d: pollen vs. male leaf. a,c: whole genome including X- and Y-specific regions, b,d: only autosomes and PAR.

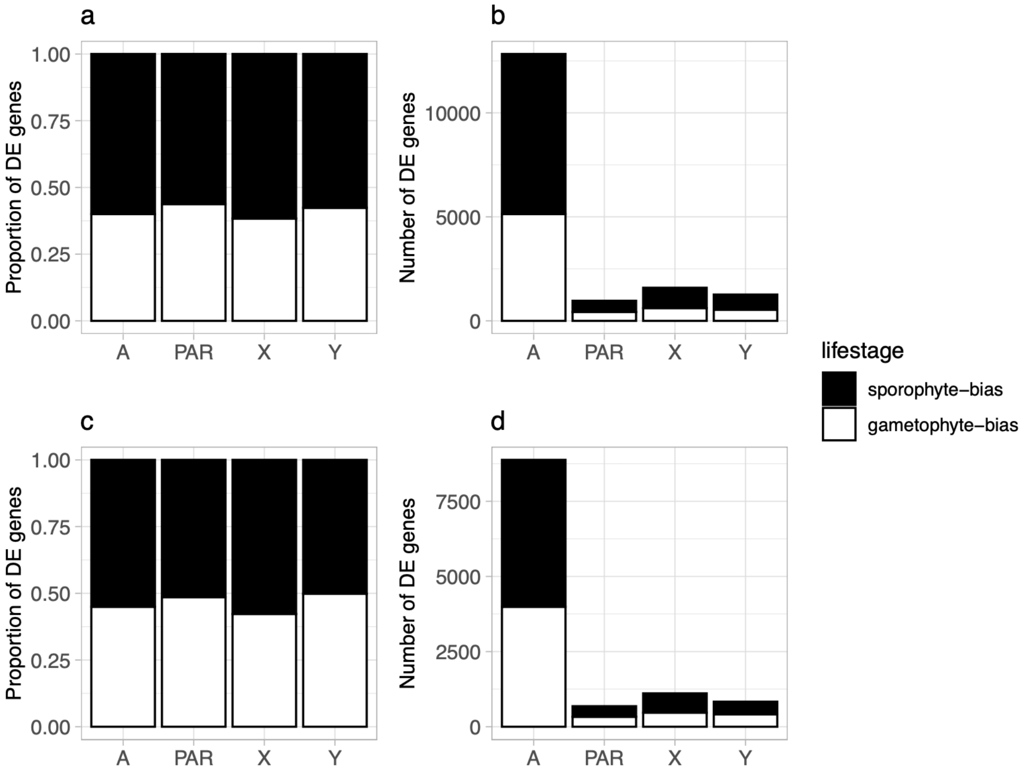


Fig S2. Enrichment of gametophyte-biased genes on the sex chromosomes in *Rumex hastatulus.* Cutoff for DE genes: adjusted $p < 0.05$ and fold change > 2 (a, b), fold change > 4 (c, d).

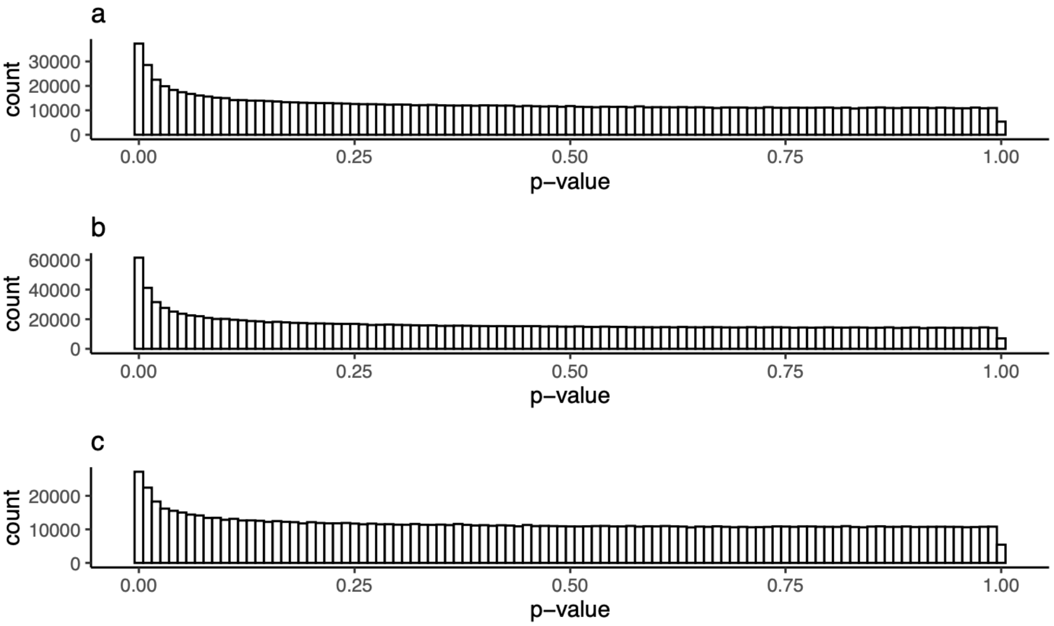


Fig S3. Distribution of nominal *p*-values for all nearby SNPs tested in male leaf (a), female leaf (b), pollen (c) of *Rumex hastatulus*. Bin size = 0.01.

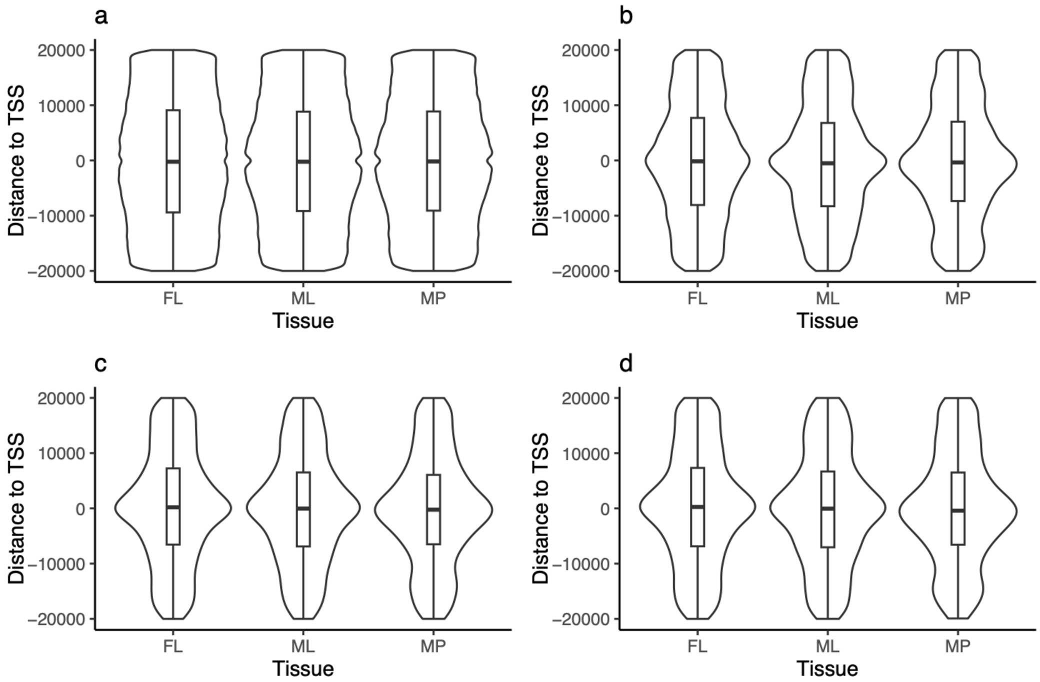


Fig S4. Distance between eQTL and the transcription start site of eQTLs male leaf (ML), female leaf (FL), pollen (MP) of *Rumex hastatulus*. a. All nearby SNPs, b. all eQTLs, c. the most significant eQTL for each eGene, d. a randomly selected significant eQTL for each eGene.

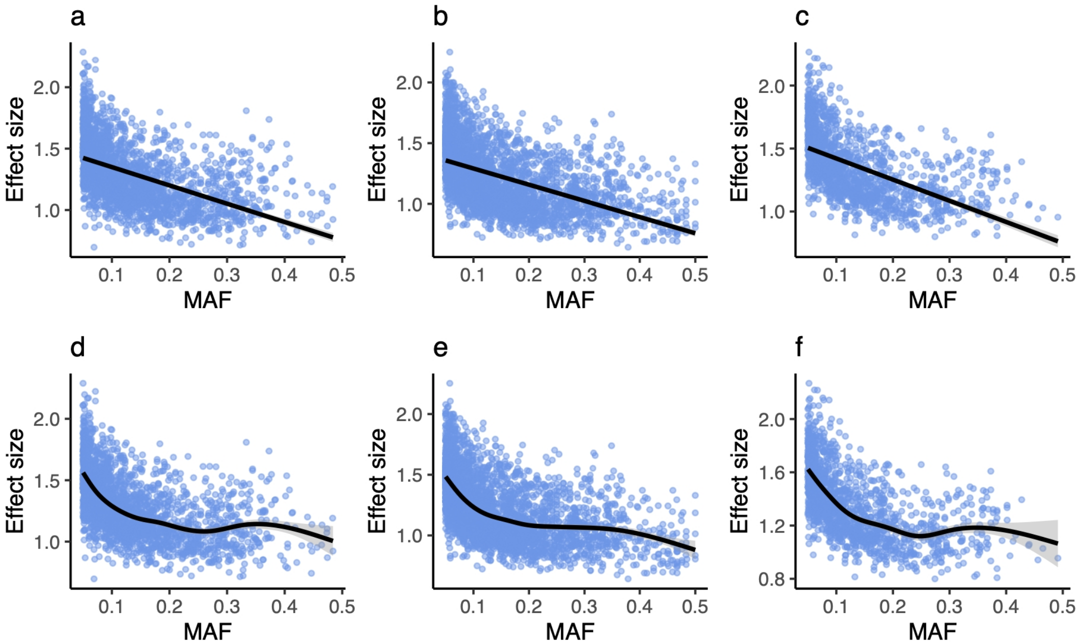


Fig S5. Correlation between effect sizes and MAFs of eQTLs in male leaf (a, d), female leaf (b, e), and pollen (c, f) of *Rumex hastatulus*. Effect size is defined as the absolute value of the slope in the linear models in eQTL mapping. a-c, linear regression, d-f: default smoothing function in R.

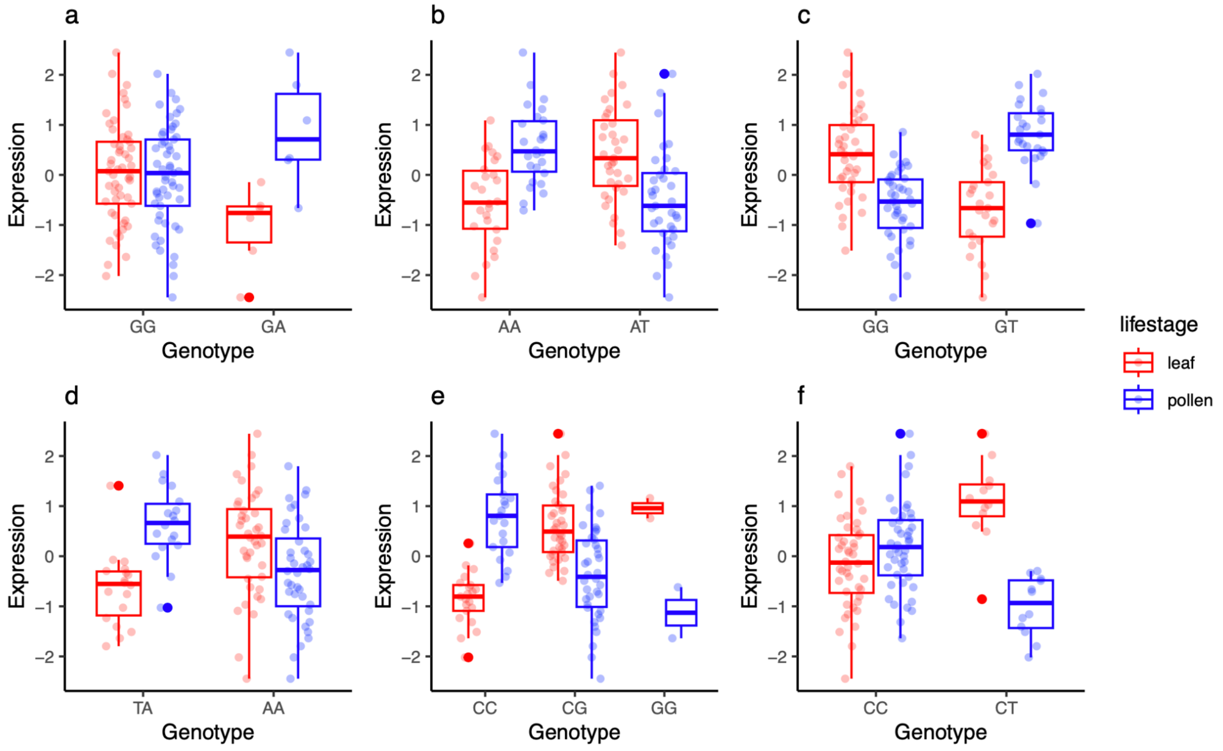


Fig S6. Examples of eQTLs showing strong discordant effects between life stages. All eQTLs are the top eQTL of the eGene in both life stages. Gene IDs and eQTL locations: a. TX_paternal_00005595 (1:255437493), b. TX_paternal_00008809 (1:455264496), c. TX_paternal_00013353 (2:162241243), d. TX_paternal_00014747 (2:238366426), e. TX_paternal_00025461 (3:13442544), f. TX_paternal_00027513 (3:75629497).

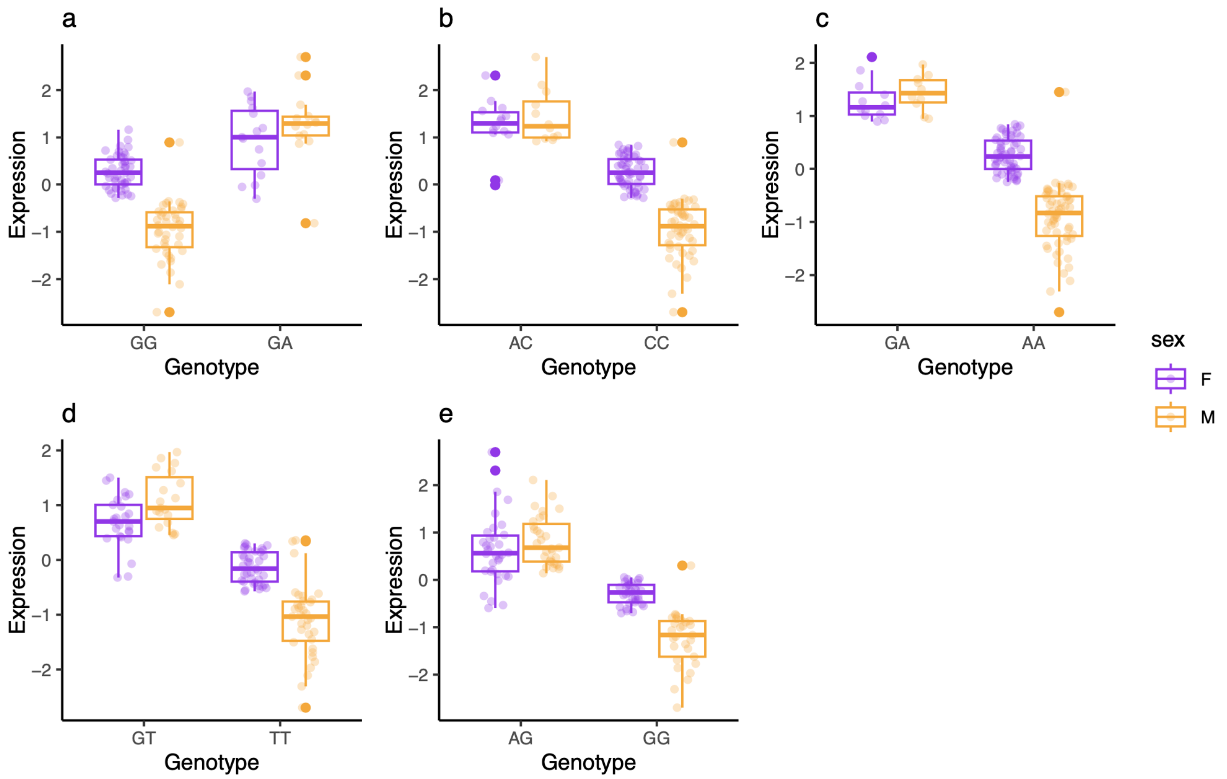


Fig S7. Examples of eQTLs showing significant Genotype × Sex interaction in *Rumex hastatulus*. Multiple-testing corrected and Benjamini-Hochberg adjusted *p*-values: 0.00197 (a), 0.00197 (b), 0.00317 (c), 0.06472 (d), 0.07921 (e). Gene IDs and eQTL locations: a. TX_paternal_00002596 (1:159008371), b. TX_paternal_00010325 (2:14033315), c. TX_paternal_00025295 (3:10654019), d. TX_paternal_00001727 (1:70371578), e. TX_paternal_00015022 (2:250749626).

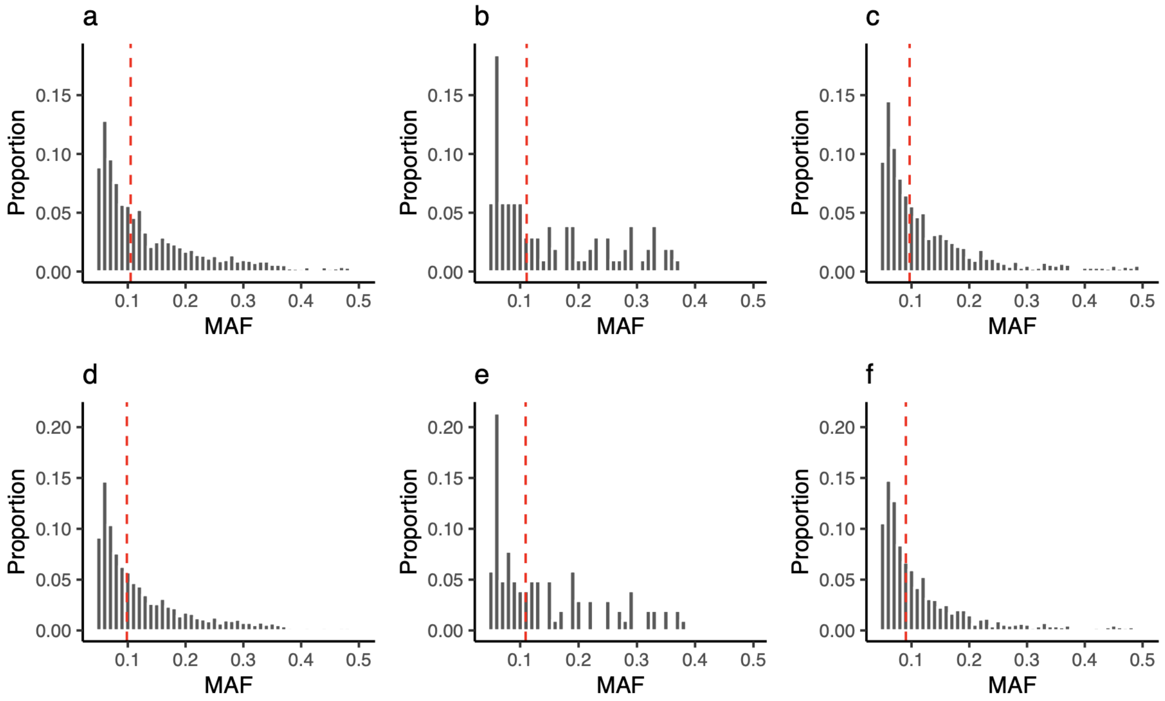


Fig S8. The distribution of MAFs of eQTLs on autosomes (a, d), PAR (b, e) and X chromosome (c, f) in female leaf of *Rumex hastatulus*. a-c: most significant eQTLs in each eGene, d-f: randomly selected eQTLs in each eGene. Binwidth = 0.01. Median of MAFs: 0.097 (a), 0.109 (b), 0.0873 (c), 0.0938 (d), 0.107 (e), 0.083 (f).

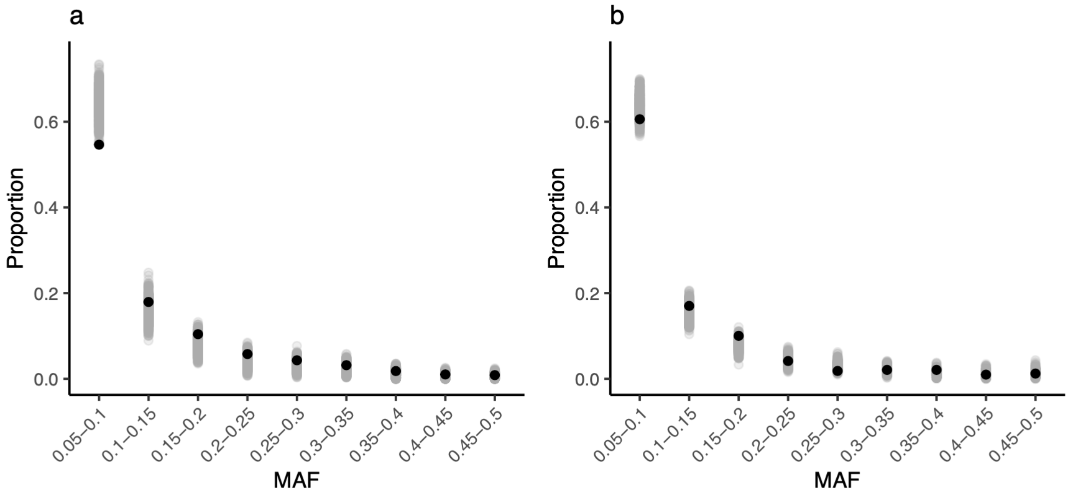


Fig S9. Distribution of MAFs of eQTLs compared to a null distribution on autosomes (a) and X-specific regions (b) in female leaf of *Rumex hastatulus*. Black dots: true positive eQTLs (observed data), grey dots: false positive eQTLs (permuted data).
